# Cognitive engagement induces area-specific fingerprints of dopamine, acetylcholine, serotonin, glutamate and GABA in prefrontal cortex and striatum

**DOI:** 10.64898/2026.05.20.726721

**Authors:** Sarah Shehreen, Seyed-Alireza Hassani, Sofia Lendor, Adam Neumann, Kanchan Sinha Roy, Janusz Pawliszyn, Thilo Womelsdorf

## Abstract

Cholinergic, dopaminergic and serotonergic neuromodulation has pervasive effects on circuit functions in prefrontal cortex (PFC) and striatum and interact with glutamatergic and GABAergic transmission. But how these neurochemicals interact during cognitive engagement is largely unknown and inferred from studying few neuromodulators at a time. Here, we sampled the extracellular availabilities of five neurochemicals in the PFC and striatum of nonhuman primates and tested how they changed when subjects switched from rest to engage in a cognitive set shifting task using miniaturized probes for diffusion-based solid-phase microextraction. Cognitive engagement was best predicted by GABAergic and cholinergic changes in the PFC, and dopaminergic and cholinergic changes in the striatum. Glutamate co-modulated with acetylcholine across states in both the PFC and striatum, while serotonin changes in PFC and striatum correlated consistent with common external modulation. These findings document an area-specific multi-neuromodulatory fingerprint of an adaptive cognitive state in the fronto-striatal network of the nonhuman primate brain.

**Teaser:** Engaging in a cognitive task reshapes neurochemical profiles across the fronto-striatal network of the primate brain

## Introduction

Cholinergic, dopaminergic and serotonergic neuromodulation exert pervasive influences on cognitive functioning across the prefrontal cortex and the striatum (Robbins and Arnsten, 2009). How each of the neuromodulator affects cognitive functions depends on the level of excitatory and inhibitory transmission in the circuit and thus varies with the availability of GABA and glutamate in the circuit (Farrant and Nusser, 2005; Sears and Hewett, 2021). Adaptive cognitive functioning of a circuit may therefore relate to the availability of multiple neuromodulators forming a complex neurochemical fingerprint (Avery and Krichmar, 2017; Shine et al., 2018; Lockhofen and Mulert, 2021).

One reason the state of multiple neuromodulators shape circuit functions are interactions among neuromodulatory systems that range from inhibition to co-transmission and co-release (Scimemi, 2025). These interactions suggests that changes in the availability of one or a few neuromodulators may change the availability of other neuromodulators. But most insights about the cross-talk of neuromodulators focus on two or three neuromodulators in a single brain area at a time. One example is striatal cholinergic receptor activation which can suppress dopamine release (Myslivecek, 2021), but may also trigger dopamine release through presynaptic acetylcholine (ACh) receptors on dopaminergic axons or indirectly when cholinergic interneurons are activated by dopaminergic-glutamatergic co-release (Soliakov and Wonnacott, 1996; Zhou et al., 2001; Threlfell et al., 2012; Scimemi, 2025). Similarly complex are interactions of striatal dopamine with GABA, glutamate or serotonin. Dopaminergic synapses in the striatum partly co-release glutamate or GABA resulting in potentially opposing effects on circuit activity (Hnasko and Edwards, 2012; Tritsch et al., 2016; Scimemi, 2025), and subsets of serotonergic receptors can trigger dopamine release (Di Matteo et al., 2008; Peters et al., 2021). These examples from the striatum raise the question whether there is a consistent pattern of available dopamine, acetylcholine, serotonin, glutamate and GABA in neural circuits that characterizes a neural circuit during efficient adaptive cognitive functioning.

Studies relating multiple neuromodulators to cognitive processes have made progress using molecular tools for real-time sensing of synaptic release, such as dLight and GRAB sensors for dopamine and acetylcholine, amongst others (Zhou et al., 2001; Muir et al., 2024; Bouabid et al., 2026). A complementary approach for relating multiple neuromodulators to cognitive processes leverages methods that evaluate their extrasynaptic concentration levels using microdialysis, cyclic voltammetry and, more recently, solid phase microextraction (Cudjoe et al., 2013; Cudjoe and Pawliszyn, 2014; Hassani et al., 2019; Lendor et al., 2019; Mayer et al., 2025). Extrasynaptic release and spillover from synaptic release concomitant with extracellular diffusion and volume transmission and has been widely documented for the major neuromodulators like dopamine and acetylcholine (Klein et al., 2019; Ztaou and Amalric, 2019; Liu et al., 2021). Extracellularly available neuromodulators curtail circuit functions by modulating synaptic activation states (Zoli et al., 1998; Liu et al., 2021; Ozcete et al., 2024). Computational approaches suggest that these neuromodulatory effects have an outsized influence on the temporal dynamics of neuronal activity (Chen et al., 2015; Shine et al., 2018; Thiele and Bellgrove, 2018), while behavioral studies have shown how varying levels of neuromodulators relate to cognition and adaptive behavior. For example, dopamine levels are linked to exploratory as opposed to exploitative behaviors (Koralek and Costa, 2021), and to the degree of motivation to engage in cognitive effortful tasks (Westbrook et al., 2020). Similarly, levels of glutamate and GABA and their ratio have major consequences for cognitive flexibility (Ajram et al., 2019). Reduced GABA levels are linked to reduced flexibility and increased repetitive behaviors (Biria et al., 2023), while altered glutamate levels are associated with deficits in multiple cognitive domains (Zhou and Danbolt, 2014). Changes in GABA and glutamate levels are associated with neuropsychiatric disorders ranging from mild-cognitive impairment to autism spectrum disorder (Waschkies et al., 2014; Marenco et al., 2018).

The surveyed studies suggest that measuring the extracellular availability of multiple neurochemicals could provide critical information to understand adaptive cognition beyond what single neuromodulator measurements can provide. Here, we make one step in this direction by leveraging principles of solid phase microextraction (SPME) to sample multiple neurochemicals simultaneously during cognitive task performance in the PFC and striatum of nonhuman primates. SPME sampling uses 200µm diameter microprobes with coated tips capable to measure dopamine, acetylcholine, serotonin, GABA and glutamate as established in in-vitro and in-vivo studies (Hassani et al., 2019; Lendor et al., 2019; Lendor et al., 2020; Bogusiewicz et al., 2021; Hassani et al., 2023). SPME sampling has comparable precision and temporal resolution to microdialysis and has major strength by sampling (i) non-disruptively by acting through passive diffusion in a way that does not deplete the free pool of neurochemicals in the system, and (ii) by using a biocompatible extracting phase that has high affinity for adsorption of multiple neurochemicals (Cudjoe et al., 2013; Cudjoe and Pawliszyn, 2014; Boyaci et al., 2020; Zhou and Pawliszyn, 2026) (see **Suppl. Text:** *Properties of SPME extraction that make it a versatile sampling procedure)*. Sampling events of 20 min. each were done twice in each experiment, at rest and while monkeys performed a set shifting task. While cholinergic and glutamatergic levels correlated in rest and task states, we found that cognitively engaging with the set shifting task was best predicted by GABAergic and cholinergic changes in the PFC, and by dopaminergic and glutamatergic changes in the striatum. While serotonin levels were unaltered by cognitive engagement, they correlated across PFC and striatum, consistent with a common external source of serotonin modulation. This result pattern shows how cognitive engagement induces area-specific neurochemical profiles encompassing glutamate, GABA, dopamine, acetylcholine and serotonin.

## Results

We sampled neurochemicals simultaneously in the PFC and the anterior striatum with SPME microprobes in 52 experimental sessions in two rhesus monkeys (subject 1: PFC n=23, striatum n=25; subject 2: PFC n=29, striatum n=27) (**Fig. 1A,B**). A first sampling event was done in the rest period (Rest state) and a second sampling event commenced while subjects engaged in an attentional set shifting task (Task state) that recruits neurons in PFC and striatum (Oemisch et al., 2019; Treuting et al., 2025) (**Fig. 1C**). The task presented in each trial three objects. Subjects chose one object by fixating it for 0.7s and received visual tokens as feedback to learn from trial-and-error which object feature was rewarded (**Fig. 1D**). The rewarded feature switched unannounced every 45-60 trials (**Fig. 1E**) requiring subjects to use attention to explore features, monitor outcomes, and update feature-value expectations using prediction errors (Womelsdorf et al., 2021).

**Fig. 1.**
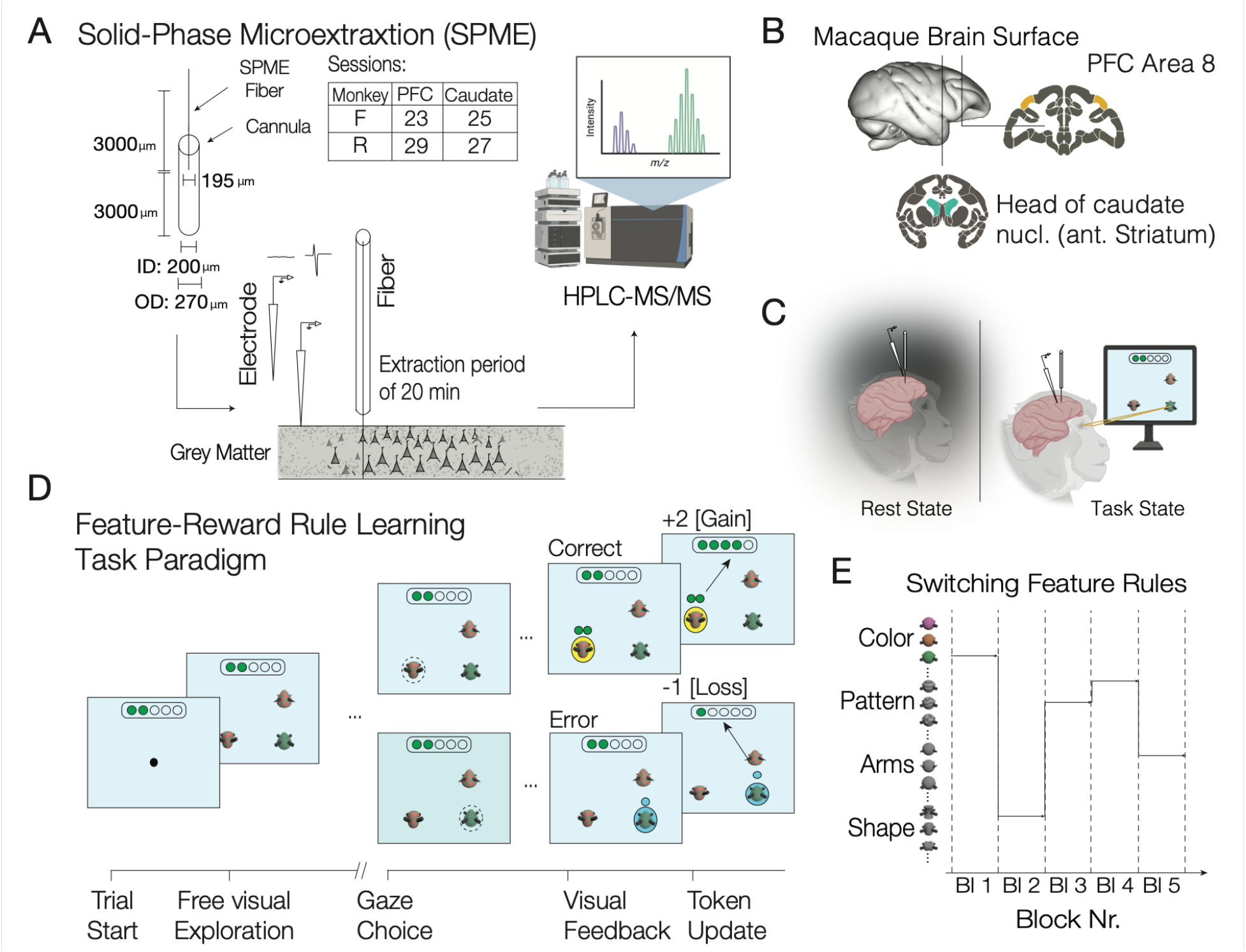
Neurochemical sampling in the prefrontal cortex and striatum during rest and task states. (**A**) A 200 µm diameter microfiber with a 2.1mm coating for Solid-Phase Microextraction (SPME) was placed in a canula, lowered to the grey matter and the coating was exposed to the tissue for 20 min. The sampled chemical concentrations were extracted as described in Methods and analyzed with mass spectroscopy. Each SPME microprobe was accompanied by a tungsten electrode used to determine the border of the white to grey matter. (**B**) Sampling sites in the prefrontal cortex and striatum of the macaque brain. (**C**) Sampling was performed during Rest and during a Task state when subjects engaged in a feature-reward rule learning task. (**D**) Task events during a trial: Three objects appeared and after a free exploration period subjects chose one objects and received feedback in the forms of tokens. (**E**) Subjects used the feedback to learn through trial-and-error which object feature was associated with reward tokens. The rewarded feature switched after blocks of 35-60 trials.

### GABA, acetylcholine and glutamate predict cognitive engagement in the PFC

Overall extracellular neurochemical concentrations in the PFC were similar in both subjects with glutamate about twice as pervasive as GABA, levels of acetylcholine lower than its precursor and metabolite choline and lower levels of serotonin (**Fig. 2A, Fig. S1A**). Extracellular levels of GABA tended to increase and cholinergic levels tended to decrease during cognitive task engagement compared to rest in absolute terms and in relation to acetylcholine, with the ratio of acetylcholine to choline increasing with task engagement (**Fig. 2B, Fig. S2**). A mixed effects logistic regression decoding model showed that the relative differences of GABA and choline distinguished Task and Rest states with 73.2% (±11.1%) accuracy (Student’s T Test, p = 0.0403) (**Fig. 2C**). The joint decoding exceeded accuracy obtained when using only GABA (69.5%, β coeff. = 3.5, p = 0.035) or only choline (58.9%, β = -1.3, n.s.) (**Fig. 2C, S3A**). Acetylcholine, serotonin, and glutamate did not allow decoding the cognitive state when used alone but predicted cognitive engagement when they were considered jointly with GABA, glutamate, or choline (**Fig. S3A**).

**Fig. 2.**
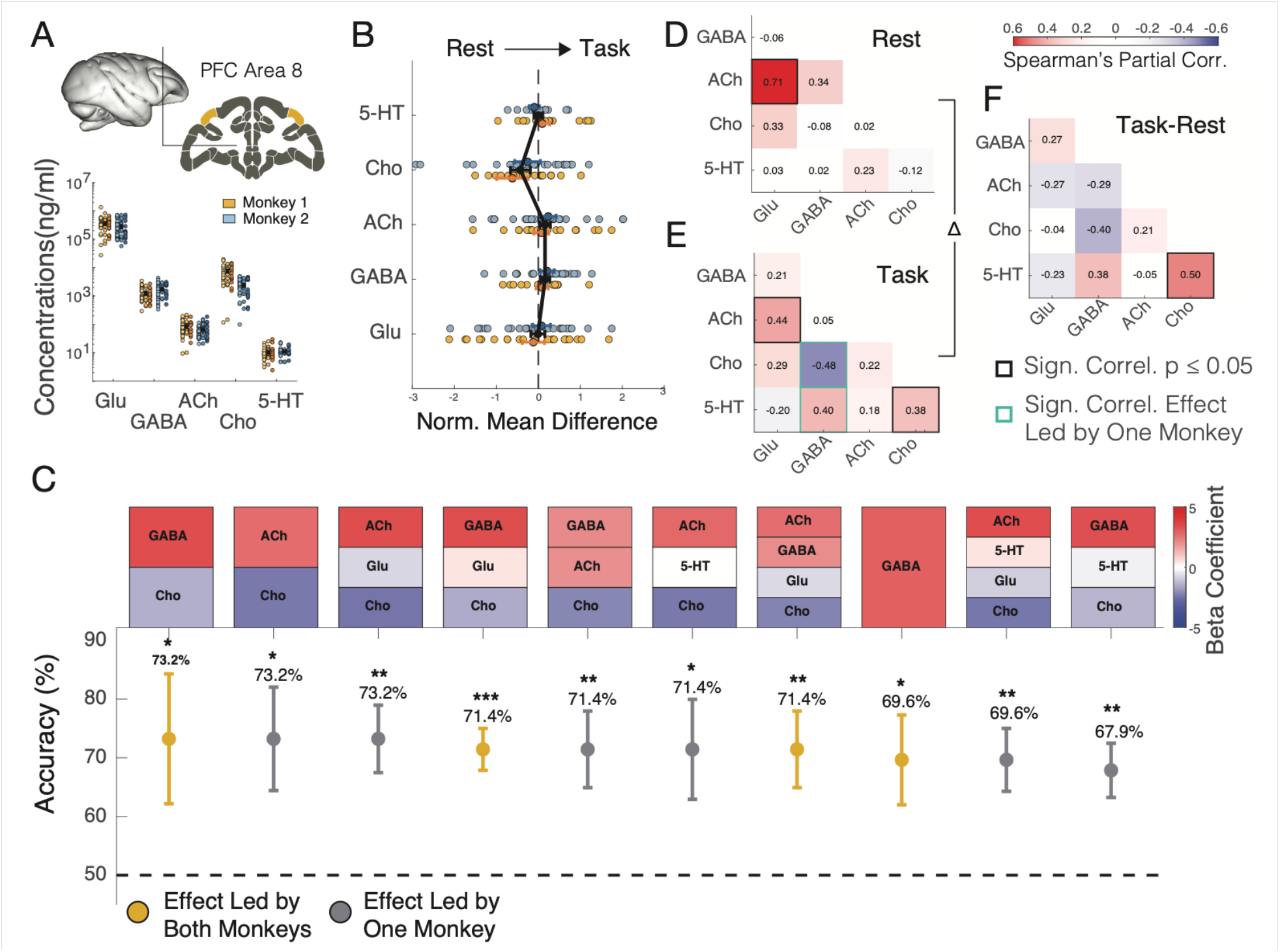
Predicting cognitive engagement versus rest state using multiple neurochemicals sampled in prefrontal cortex (PFC). (**A**) Neurochemical concentrations in the PFC for monkey 1 (yellow) and 2 (blue). Lighter colors are from Rest state, darker from task states. (**B**) Mean differences (Task minus Rest state) of session normalized concentrations of all neurochemicals. (**C**) Accuracy of the ten combinations of neurochemicals that best classify Task vs Rest state (best ranked on the left) using a generalized non-liner mixed effects model with 7-fold cross-validation. The top row shows the color-coded beta-coefficients of the neurochemicals underlying the accuracy shown in the bottom panel. Red colors denote neurochemicals predicting the task state with increased concentrations relative to the rest state; blue denotes neurochemicals reduced in the task state. See Fig. S3 for neurochemical concentrations with accuracy ranked eleventh onwards. (**D**,**E**) Correlations of neurochemical concentrations simultaneously sampled during in the Rest State (*D*) and the Task state (*E*). (**F**) Difference of concentrations in the Task minus Rest state. Cells with thick black border in *D-F* denote p<0.05 in both subject; green borders denote p<0.05 effect evident in only one of the subjects. Error bars are SE’s.

The decoding results suggest that neurochemicals were not independently varying in the PFC. Correlation analysis confirmed that glutamate and acetylcholine correlated in the PFC in the Rest and Task state (Spearman’s partial rank correlation, FDR corrected, Rest: r = 0.71, p = 0.001; Task: r = 0.439, p = 0.0265) (**Fig. 2D,E**; **Fig. S4A**,**B**). In addition, while choline and serotonin were unrelated in the rest state, they were correlated during cognitive engagement (Spearman’s partial rank correlation: r=0.378, p=0.05) (**Fig. S4C**). The choline-serotonin correlation increased between states (difference Task-Rest, permutation testing, p = 0.046) (**Fig. 2E**). To test whether the serotonin-choline co-modulation was linked to cognitive demand we separated sessions in which set shifting was performed with simple objects (varying in one feature: low interference) or with more complex objects (varying in features of 2 dimensions: higher interference). We found that the choline-serotonin correlation increased specifically during performance of the high-interference task compared to the low-interference task (difference in correlation, permutation testing: High-interference task vs. Rest: 0.64, p<0.05; Low-interference task vs. Rest: 0.13, n.s.) (**Fig. S5**).

Neurochemical changes during the Task state might be influenced by the number of earned fluid rewards, the speed of choice reaction times, the overall performance levels or how fast subjects learned the new object sets after a set shift. Analyzing these factors showed that in the PFC no systematic correlations of these performance markers for glutamate, GABA, acetylcholine and serotonin, and non-systematic changes of choline concentrations with overall performance and reward rate (**Fig. S6B-C**). Similarly, neurochemical levels were not linked to physiological changes in heart rate of blood oxygen levels (**Fig. S7**).

### Dopamine, glutamate and acetylcholine predict cognitive engagement in anterior striatum

We next analyzed neurochemical concentrations in the anterior striatum (head of the caudate nucleus). Subjects showed similar levels of glutamate, GABA, acetylcholine, choline and serotonin (**Fig. 3A**). In addition, dopamine levels were robustly measurable in the striatum at concentrations similar to those of acetylcholine (**Fig. 3A**). Compared to the PFC, striatal glutamate levels were lower and GABA levels higher (**Fig. S1**).

**Fig. 3.**
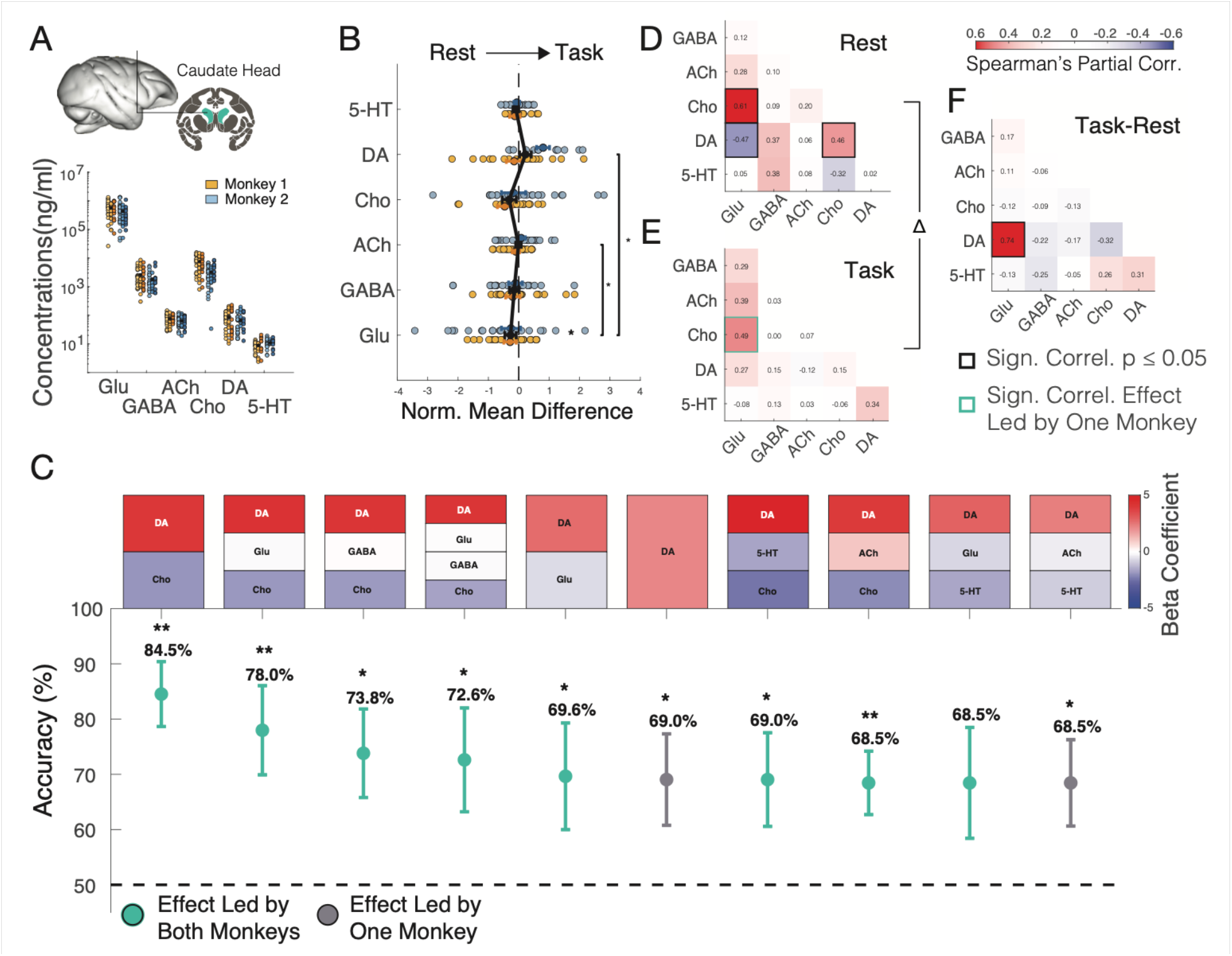
Predicting cognitive engagement versus rest state using multiple neurochemicals sampled in the anterior striatum (head of the caudate nucleus). (**A-F**) Same as Fig. 2 A-F but for neurochemicals sampled in the striatum.

Striatal dopamine concentrations increased during cognitive task engagement compared to the rest state relative to glutamate (Kruskal-Wallis Test with Wilcoxon’s for pairwise test with FDR correction, p = 0.0102) (**Fig. 3B**). Similarly, acetylcholine levels moderately increased during the task versus rest state relative to glutamate levels (Kruskal-Wallis Test with Wilcoxon’s for pairwise test with FDR correction, p = 0.0415) (**Fig. 3B**), with a trend for a higher ratio of ACh to choline during the task than rest state (p = 0.0885, **Fig. S2**). Consistent with these changes, dopamine was the best single predictor of cognitive engagement (accuracy: 69% ± 4.9%, Student’s T Test, p = 0.0027), with increased decodability of the task state when dopamine was considered jointly with choline (accuracy: 84.5% ± 9.2%, Student’s T Test, p = 0.0016). This joint decoding was based on dopamine increases (β = 4.1763) going along with choline decreases (β = -1.9) in the task state compared to the rest state (**Fig. 3C, Fig. S3B**). The task state was also jointly decodable using dopamine and choline together with glutamate (accuracy: 78%, Student’s T Test, p<0.01), GABA (accuracy: 73.8 %, Student’s T Test, p<0.05), serotonin (accuracy: 69%, Student’s T Test, p<0.05), and acetylcholine (accuracy: 68.5 %, Student’s T Test, p<0.05) (**Fig. 3C**). These results illustrate the particular role of dopamine to index task states and that dopamine’s availability is linked to the modulation of the other measured neurochemicals (*see* discussion, **Fig. S3B**).

We next quantified the co-variation of neurochemical concentrations in the striatum. Similar to the PFC, cholinergic concentrations positively correlated with glutamate in the striatum both at rest and during the task state (Spearman’s partial rank correlation, FDR corrected, Rest: r=0.61, p=0.003; Task: r=0.488, p=0.016) (**Fig. 3D-F, Fig. S8A**). In contrast, dopamine correlated with glutamate negatively during rest (Spearman’s partial rank correlation, FDR corrected, r = -0.465, p = 0.0349), but positively during the task state (dopamine – glutamate difference task vs rest state, permutation test, p = 0.006) (**Fig. 3F**). The dopamine-glutamate co-modulation during the task state was more pronounced in sessions in which the set shifting task used more complex visual objects, indicating it dependent on cognitive demands (dopamine – glutamate correlation difference task vs rest state, permutation test, r diff: 1.045, p=0.002) (**Fig. S8C-F**). Control analysis supports this suggestion by showing that the total number of obtained rewards, the reward rate, or other performance measures did not correlate with dopamine, glutamate, GABA, or acetylcholine levels (**Fig. S6D**,**E**). Similarly, heart rate variations were not related to variations in neurochemical levels with the exception of higher acetylcholine levels associated with lower heart rate and oxygen saturation (ACh x heart rate: r = -0.475, p = 0.0059; oxygen levels: r = -0.364, p = 0.0268) (**Fig. S7B**).

### PFC-Striatum co-modulation of serotonin and anti-correlation of GABA and acetylcholin

In 50 experimental sessions (subject 1 / 2: n = 23 / 27) neurochemicals were simultaneously sampled in the PFC and the striatum allowing to test whether neurochemical concentrations in the PFC predicted changes in the striatum. First, we found serotonin levels in the PFC and the striatum were positively correlated during rest and task states (Rest: r = 0.52, p < 0.0132; Task: r=0.46, p < 0.0189) (**Fig. 4A-D**). This finding is consistent with serotonin levels in the PFC and the striatum originating from a similar external source.

**Fig. 4.**
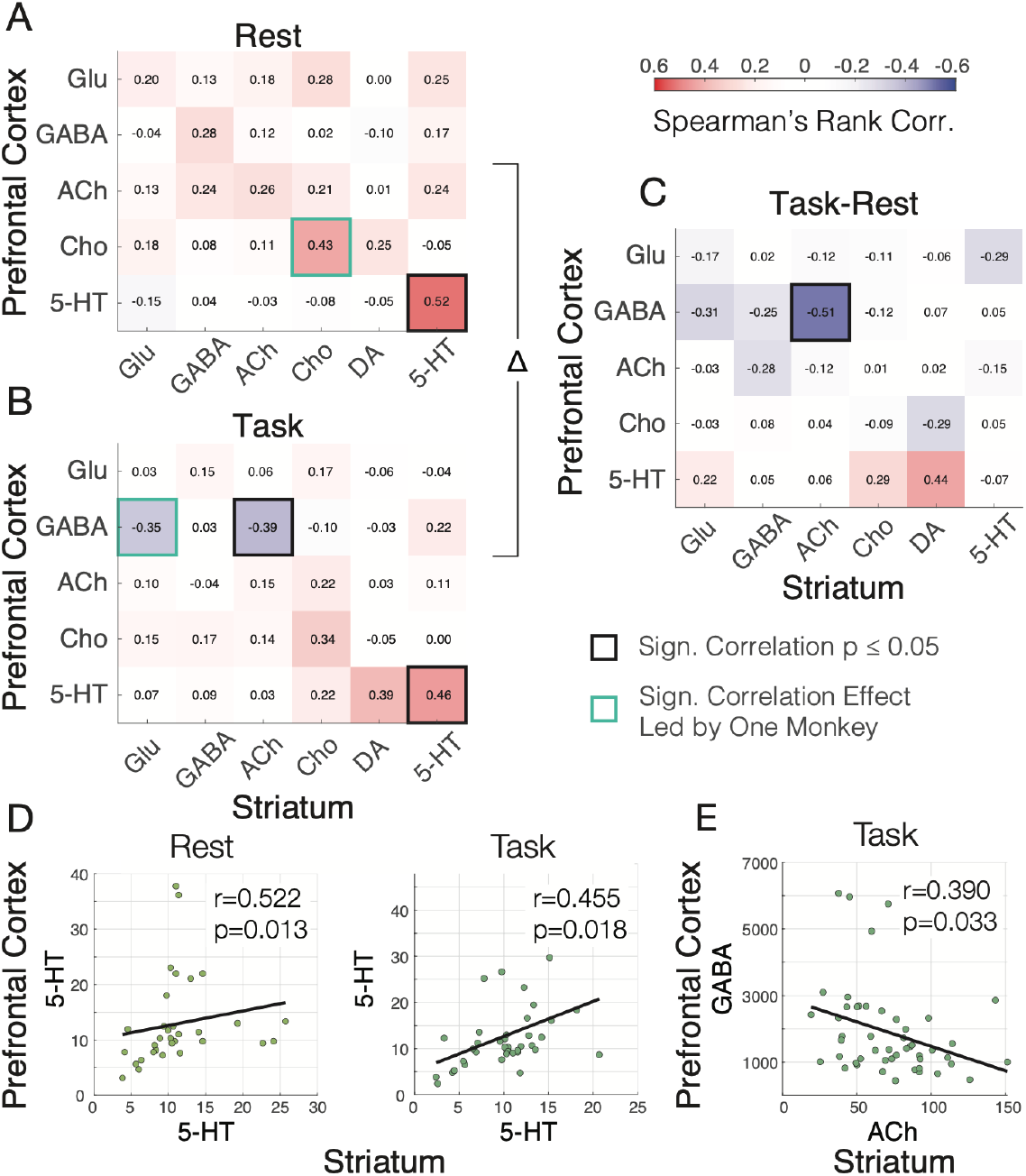
Co-modulation of prefrontal cortex and striatum neurochemical changes at rest and during cognitive engagement. (**A**,**B**) Correlation of neurochemical changes in the PFC (*y-axis*) and the anterior striatum (*x-axis*) during the rest state (*A*) and during cognitive task engagement (*B*). Thick outlined cells denote p<0.05 as evident in both subjects (black) or based on one subject (green). (**C**) Correlation difference of [Task minus Rest] state neurochemical levels. PFC GABA levels correlated with striatum ACh levels (p<0.05). (**D**) Correlation of serotonin levels of the PFC (*y-axis*) and striatum (*x-axis*) at rest (*left*) and during task engagement (*right*). (**E**) Correlation of GABA levels of the PFC (*y-axis*) and ACh level of the striatum during the task state.

Second, we found that PFC GABA levels and striatal cholinergic levels showed a task dependent anti-correlation (difference Task versus Rest permutation test: r = -0.51, p = 0.023) (**Fig. 4A-C**). PFC GABA was uncorrelated to striatal acetylcholine in the rest state (r = 0.12, n.s), but it was anti-correlated in the task state (r = -0.39, p = 0.033) (**Fig. 4E**). The association of increased GABA levels in the PFC with lower cholinergic availability in the striatum was more pronounced when the task involved high interference control (r = -0.63, p = 0.0150), but remained negative also during performance of the task condition with lower interference control demand (r = -0.35, n.s: p = 0.2260).

## Discussion

Here, we documented a rich pattern of neurochemical co-modulation in the PFC and striatum of nonhuman primates and how this pattern changed when subject engaged in a cognitively demanding attentional set shifting task. First, irrespective of the cognitive task cholinergic and glutamatergic concentration levels were positively correlated in both, the PFC and the striatum (**Fig. 2D-F, 3D-F**). Secondly, cognitive task engagement changed individual neurochemical concentrations and their co-modulation systematically. In the PFC GABAergic and cholinergic changes maximally predicted task engagement, reaching 73.2% state decoding accuracy (**Fig. 2C**), while in the striatum dopamine increases were most predictive of cognitive task engagement together with cholinergic and glutamatergic modulation, reaching 84.5% and 78% state decoding accuracy, respectively (**Fig. 3C**). A third pattern of result was evident for serotonin levels, which correlated in PFC and striatum (**Fig. 4**), but showed increased co-modulation with cholinergic levels during cognitive engagement only in the PFC (**Fig. 2D-F**).

### Widespread co-modulation of glutamate and acetylcholine in PFC and striatum

Glutamate and acetylcholine positively correlated in the PFC (Rest / Task: r = 0.71 / 0.439) and in the striatum (Rest / Task: r=0.61 / r=0.488) at rest and during task engagement. This widespread correlation is consistent with cholinergic neurons co-releasing glutamate (Prado et al., 2017). The lack of task modulation suggests that extracellular glutamate levels did not accumulate over the 20 min sampling period when PFC and striatum circuits are more active during set shifting. Accumulation of glutamate has been observed with more excessive cognitive effort (Wiehler et al., 2022), reflecting a state when subjects are overwhelmed by task demands (Zhou and Danbolt, 2014). Our pattern of results suggest that such detrimental task demands were not reached with our set shifting task. The task condition with higher demands using more complex objects was performed well above the 33% chance level at ∼65% overall accuracy during the 20 min. sampling period (**Fig. S6A**). For this more demanding task, glutamate increased co-modulation with GABA (task versus rest difference of correlation: 0.64, permutation test; p=0.0430) (**Fig S5C**), raising the possibility that balancing glutamate and GABA levels indexes successful coping with cognitive demands.

### Prefrontal cortex GABA and ACh/choline ratio predict cognitive engagement

In the PFC cognitive engagement was best predicted by GABA level increases (69.5% accuracy) which, when combined with reduced choline distinguished task and rest states at 73.2% accuracy (**Fig. 2C**). This finding documents the importance of PFC GABA levels to support adaptive cognition and is consistent with the detrimental effects of GABA antagonists (e.g. bicuculline) to impair working memory performance in NHPs (Sawaguchi et al., 1989), and impair set shifting and increase perseveration in rodents (Enomoto et al., 2011). Genetic studies likewise suggest that reduced GABA levels are linked to lower recruitment levels of the PFC and concomitant more stereotyped repetitive behaviors (Ajram et al., 2019; Wang and Sun, 2025). Our findings extend these insights by documenting that an increase in GABA levels in the PFC is predictive of successful cognitive task engagement.

The predictive value of GABA was increased when the decoder used it jointly with choline. While this analysis revealed that task engagement is accompanied by moderate decreases of choline, a more complete picture takes into account that choline is a precursor and metabolite of ACh and may not independently modulate across states (**Fig. 2C**). We confirmed this dependency by observing that the ratio of prefrontal ACh/Cho systematically increased in the task versus the rest state (Wilcoxon’s test, p = 0.003) (**Fig. S2**). This finding shows that cognitive engagement changed the balance of ACh to choline, while ACh and choline on their own show only moderate effects across the analyses. This finding contrasts to a hypothesized reciprocal relationship of ACh and choline (Koshimura et al., 1990; Ikarashi et al., 1997), and rather supports earlier suggestions that the relationship of ACh and choline is complex and relatively weak, with choline modulation by itself is a relatively poor predictor of cholinergic transmission (Parikh et al., 2004; Sarter and Parikh, 2005) (*see* also **Suppl. Text:** *Relationship of acetylcholine and choline levels in striatal and cortical circuits*).

### Striatal dopamine predicts cognitive engagement together with glutamate

Dopamine was the single most predictive neurochemical predicting the task state in the striatum. This finding is consistent with prior studies showing how elevated dopamine promotes attentional control and cognitive effort (Krebs et al., 2012; Westbrook and Braver, 2016; Westbrook et al., 2020) and scaling with reward prediction errors and the behavioral propensity to choose the more rewarding option in a decision making task (Pessiglione et al., 2006). Dopamine indexed cognitive engagement not in isolation. We found that striatal dopamine and glutamate concentrations were anticorrelated at rest and shifted to a positive correlation when subjects engaged in the cognitive task (**Fig. 3D-F**). This finding likely involves direct co-release processes as well as indirect co-modulation from increased cortico-striatal glutamatergic synaptic activity and local dopamine release. Striatal co-release of dopamine and glutamate has been documented in a subset of dopaminergic synapses (Chuhma et al., 2014; Scimemi, 2025) and striatal D1 receptor activation has been shown to enhance AMPA and NMDA receptors kinetics and increases the response to glutamate (Hallett et al., 2006; Surmeier et al., 2007). Our findings resonate with these reports showing that successful learning in NHP’s is accompanied by concomitant increases of dopamine and glutamate.

### Tonic cholinergic interactions with dopamine in the striatum

While increases of striatal dopamine levels predicted the task state at 69% accuracy, the predictive power increased to 84.5% when the decoding model additionally considered variations of choline (**Fig. 3B**). This finding reflects that dopamine and choline independently contributed to predicting the task state. Choline levels were lower during the task state (reflected in a negative beta weight), while levels of ACh moderately increased during the task state (reflected in a positive beta weight). Similar to the PFC, the net ratio of ACh to choline showed a trend to be positive (p = 0.0885, **Fig. S2**). These observations suggest a net positive association of moderate increases in ACh relative to choline during more active states (see also **Suppl. Text**: *Relationship of acetylcholine and choline levels in striatal and cortical circuits*). This interpretation resonates with previous evidence of a net positive association of cholinergic activation triggering dopamine release through presynaptic ACh receptors on dopaminergic axons, as well as dopaminergic-glutamatergic co-release triggering cholinergic interneuron activity (Soliakov and Wonnacott, 1996; Zhou et al., 2001; Threlfell et al., 2012; Scimemi, 2025). Beyond these positive ACh-dopamine interactions over slower 20min. sampling time scales, there is, however, also evidence for transient anticorrelations of dopamine and ACh during reward processing (linked to increases in dopamine) and action initiation (linked to increases in ACh) (Chantranupong et al., 2023; Bouabid et al., 2026). Together these findings point to the importance of evaluating ACh-dopamine interactions at multiple timescales.

### Serotonin co-modulation across PFC and striatum

While serotonin levels did not distinguish the task state from rest in either brain area (**Fig. S3**), they correlated across the PFC and the striatum at rest (r=0.52) and during task engagement (r=0.46) (**Fig. 4**). This widespread co-modulation suggests a common source consistent with serotonin projections from the dorsal raphe nucleus having broad and heterogeneous projections encompassing PFC and the striatum (Ren et al., 2019). The absent task modulation of serotonin suggests that neural firing changes in the dorsal raphe nucleus that are related to the anticipation of expected rewards or the processing of reward outcomes (Zhong et al., 2017; Spring and Nautiyal, 2024) are not translated into tonic increases of serotonin levels in the PFC and striatum. There is substantial heterogeneity of dorsal raphe neuron firing during behavior (Okaty et al., 2019), suggesting that finer grained behavioral assays and measurements of direct synaptic transmission will be necessary to understand how serotonin contributes to striatal and PFC functions.

### Multi-neuromodulator fingerprint of cognitive engagement across the frontostriatal network

Our study documents how multiple neurochemicals are recruited during cognitive engagement. Prominent changes of GABA in the PFC and dopamine in the striatum most apparently predicted the transition from a rest to an active task processing state. In both brain areas, cognitive engagement triggered interactions of GABA and dopamine with cholinergic, glutamatergic and serotonergic levels, showcasing the multi-neuromodulator fingerprint of cognitive engagement in the nonhuman primate fronto-striatal network. The emerging view is a tonic pattern of extracellular available neurochemicals that will curtail neuronal processing during adaptive cognitive behavior. How the extracellular availability of neurochemicals translate into adaptive neuronal processing will require combining the measuring tonic levels of neurochemicals and more transient and spatially specific synaptic release processes while subjects engage in cognitive task performance (Zhou et al., 2001; Muir et al., 2024; Bouabid et al., 2026).

## Materials and Methods

All animal care and experimental protocols were in accordance with the National Institutes of Health Guide for the Care and Use of Laboratory Animals, the Society for Neuroscience Guidelines and Policies, and approved by the Vanderbilt University Institutional Animal Care and Use Committee. Data was collected from two pair-housed adult male rhesus macaques (Macaca mulatta; age: 8 and 10 years-old; weight: 12 and 14 kg). Both monkeys were under fluid control and worked at the same time every day to acquire at least 20 ml/kg/day of water.

### Experimental Design

#### Experimental setup

The experimental setup was similar to a previous study (Hassani et al., 2023). Subjects were seated in a chair and placed in a custom-built sound attenuated booth in front of a 21’’ LCD screen 63 cm from the animals’ eyes, where they completed a behavioral task for water reward. Behavior, visual display, stimulus timing, and reward delivery were controlled by the Unified Suite for Experiments (USE), which integrates an IO-controller board with a unity3D video-engine based control for displaying visual stimuli, controlling behavioral responses, and triggering reward delivery (Watson et al., 2019). Visual stimuli were 3D-rendered multi-dimensional so-called Quaddle objects (Wen et al., 2025) presented at random locations with 4.1’’ eccentricity from the center of the screen. The objects were ∼1.1’’ in diameter and varied across one or two feature dimensions (body shape, color, pattern or arm shape/angle) with the remaining feature dimensions being identical between objects per block.

Each subject was implanted with a custom build titanium headpost and a 22 mm x 36 mm recording chamber (Rogue Research, Montreal, CA) over the frontal region of the left hemisphere guided by stereotaxic coordinates (Paxinos et al., 2000) and magnetic resonance (MR) images. The trajectories and depth for the neurochemical sampling were determined using CT images of the skull after the chamber was implanted, co-registered to the MR images obtained prior to the implantation. Anatomical reconstruction and segmentation of brain areas were done as in previous studies (Treuting et al., 2025).

#### Solid-phase microextraction sampling probe and quantitation of neurochemicals

Neurochemicals were extracted from the prefrontal cortex and striatum using miniaturized microprobes built and optimized for neuromodulator sampling based on principles of Solid-Phase Microextraction (SPME) (Pawliszyn, 2000, 2012). The design, fabrication and optimization of the probes are described in detail elsewhere (Lendor et al., 2019). Each SPME fiber consisted of a 200 µm stainless steel wire which was coated at the tip over a length of 3 mm as described in (Lendor et al., 2019) and validated in the nonhuman brain in (Hassani et al., 2019). The coating served as the extracting phase and consisted of an in-house synthesized hydrophilic-lipophilic balance polymer functionalized with benzenesulfonic acid to introduce strong cation exchange properties (HLB-SCX). Prior to coating the tip was acid-etched down to an approx. 100 μm diameter so that the deposition of a 50 µm layer of extracting phase particles resulted in the same 200 µm diameter as the shaft of the probe.

#### Experimental procedure

During each experiment, animals were seated in the sound attenuating booth and prepared for simultaneously sampling of neurochemicals with SPME probes (*below*) and electrophysiological recording of spiking activity using tungsten electrodes (**Fig. 1A**). An experiment began by lowering tungsten electrodes to the grey matter in the prefrontal cortex (area 8) and the anterior striatum (head of the caudate nucleus) (**Fig. 1B**) in order to demarcate the depth at which neuronal spiking activity signaled the begin of grey matter. Following the demarcation of grey matter, two sets of neurochemical measurements were taken using biocompatible microprobes operating under the principles of solid phase micro-extraction (SPME) (Pawliszyn, 2000). The first measurement was taken while the subject was at rest and before a behavioral task began, while the second measurement was taken once subjects had started to engage with the behavioral task (Rest and Task state in **Fig. 1C**). In a subset of experimental sessions, the first and second measurements were done while the subjects performed different tasks which varied in the number of irrelevant, distracting visual features that varied from trial-to-trial (task conditions: low and high distractor interference).

#### SPME neurochemical sampling and electrophysiological recording

Before each experiment the SPME probes were cleaned with the mixture of methanol, acetonitrile and 2-propanol (50:25:25, v/v/v), activated in the mixture of methanol and water (50:50. v/v) and sterilized in steam for 15 min at 121°C. During the sampling, SPME probes were inserted into sheathing cannulas to protect the coating and prevent it from extracting compounds on the way to the target brain area (**Fig. 1A**). The cannulas underwent the same cleaning and sterilization procedure as the probes and then the lengths of both components of the SPME assembly were adjusted to 60-70 and 70-80 mm for the probes and cannulas, respectively. Each SPME-probe and respective cannula were lowered into the brain using software-controlled precision micro-drives (Neuronitek, Ontario, Canada) alongside an accompanying tungsten electrode (FHC). The tungsten electrode was mounted ∼1mm next to the SPME probe and cannula and its depth was controlled by a separate micro-drive. The electrode was lowered first to identify the depth at which spiking activity of neurons was evident marking the start of grey matter in the dorsolateral prefrontal cortex (PFC, area 8) and the anterior striatum (head of caudate nucleus). The cannula with the shielded SPME probe was lowered a few micrometer below the start of the grey matter before the SPME probe was driven out and exposed to gray matter for 20 min before being driven back into the cannulas and pulled back out of the brain (**Fig. 1A**). Once outside of the brain, SPME probes were rinsed in ultra-pure water and any visible tissue removed before suspending the probes in a glass vial and subsequently freezing them in a -80 °C freezer.

During each experimental session two SPME sampling events took place (each with one probe in the PFC and one probe in the striatum), which was accomplished by switching the two multi-drive assemblies that held the SPME probe, cannulas, and tungsten electrode. Prior to the experiment, four of these assemblies were prepared. During the experiment, the first two assemblies were used for the first sampling event and the second pair of assemblies were used for the second sampling event. Notably, each pair of assemblies coupled to a stationary positioning system such that the same locations were being sampled by the two sampling events per day. Switching the assemblies took ∼5 min. After the experiment, samples were stored in the -80 °C freezer for up to 2 weeks before being transported in a dry-ice box for desorption and quantitation. Reconstruction of SPME sampling locations were done in our previous study (Hassani et al., 2019).

#### Determination of the 20 min sampling duration

An extraction/sampling duration of 20 min was selected to ensure sampling provided sufficient recoveries and good reproducibility of measurements, while maintaining temporal resolution (**Suppl. Text**: *Properties of SPME extraction that make it a versatile sampling procedure*). The kinetics of target analyte extraction has been visualized in vitro through extraction time profiles in (Lendor et al., 2019). In summary, brain homogenate was spiked with known concentrations of target analytes and four replicates per time point were collected during which multiple SPME probes were placed in the homogenate and sequentially removed at progressive times after initial insertion and quantified. The resulting extraction time profiles document the extracted amount of analyte over time. Each analyte proceeds through a linear “kinetic” regime, a “dynamic” regime and an “equilibrium” regime over the extraction time. These labels describe the various rates of analyte adsorption on the SPME probe, where the kinetic regime reflects the fastest uptake of the target analyte (typically within 1-5 min across neurochemicals), and the final equilibrium regime reflects the period of only marginal further analyte collection, in this case between 20-45 min depending on the particular neurotransmitter (Lendor et al., 2019). The 20 min extraction duration was chosen to balance good sensitivity and precision for multiple neurochemicals with reasonable temporal resolution of measurements. It is noteworthy that the extraction kinetics are independent of the concentration of the analyte in the extracellular milieu (Kasperkiewicz et al., 2023).

#### Neuromodulator detection and quantitation

All chemicals, reagents, and materials used for chemical analysis are described in the **Suppl. Text:** *Chemicals, reagents and materials*. The post-processing included LC-MS/MS analysis, and detailed quantitation of neuromodulators is described in the **Suppl. Text:** *Quantitation of neuromodulators*. In brief, the SPME probes were desorbed into 50 µL of acetonitrile/methanol/water 40:30:30 solution containing 0.1% formic acid and internal standards at 20 ng/mL for 1 h with agitation at 1500 rpm. LC-MS/MS analysis was conducted to quantify dopamine, choline and glutamate. Details of the instrumental LC-MS/MS analysis with and without derivatization of target neurochemicals, are included in **Table S1**. 20 µL of each remaining extract was subsequently transferred into an insert vial and evaporated to dryness at 30°C in order to complete derivatization with benzoyl chloride, according to a refined procedure described by Song et al. and Wong et al. (Song et al., 2012; Wong et al., 2016). 10µL of 50 mM sodium tetraborate buffer (pH 9.2) was added to the dry residue, followed by vortexing and addition of 10 µL of benzoyl chloride solution (2% in acetonitrile, (*v/v*) and vortex mixed again). Then, LC-MS/MS analysis was conducted to quantify serotonin, acetylcholine and GABA (details in **Suppl. Text**: *Post-desorption derivatization with benzoyl chloride*). The LC-MS/MS analysis was carried out using an Ultimate 3000RS HPLC system coupled to a TSQ Quantiva triple quadrupole mass spectrometer (Thermo Fisher Scientific, San Jose, CA, USA). Data acquisition and processing were performed using Xcalibur 4.0 and Trace Finder 3.3 software (Thermo Fisher Scientific, San Jose, CA, USA).

#### MRI guided electrophysiological mapping of target tissue

The anatomical coordinates of the brain regions of interest were first identified through 3 T MR images. The MR images were then verified with extracellular electrophysiological recordings of the target areas, which provided the gray and white matter boundaries for the cortical sites and the dorsal most aspect of the head of the caudate nucleus. Tungsten microelectrodes were 200 μm thick with an impedance of 1-2 MΩ. All electrodes, SPME probes and their accompanying guiding cannulas were driven down into the brain and later out using software-controlled precision microdrives (Neuronitek, ON, Canada). Electrodes were connected to an Intan Recording System (RHD2000, Intan Technologies, LLC) and wideband and spiking data visualized during the recording using Open-Ephys (Siegle et al., 2017) (open-ephys.org). For every recording day, electrodes were lowered until the first detection of spiking activity (indicative of gray matter) at the depth suggested by the MR images.

#### Behavioral task paradigm

The behavioral task was a feature-reward rule learning task described in previous studies (Womelsdorf et al., 2021; Hassani et al., 2024). Subjects had to learn which visual object feature was associated with reward feedback within blocks of 45-60 trials (**Fig. 1D,E**). A trial started when subjects fixated on a central point for 500 ms, which was followed by the presentation of three objects. Subjects could freely view those objects and choose one object to receive feedback by fixating an object for 0.7 s. Only one object had a feature that was associated with reward. Choosing the object with the rewarded feature resulted in visual (yellow halo) and auditory (high pitch tone) feedback and fluid reward while choosing an object without the rewarded feature resulted in a grey halo and low pitch tone feedback an no reward. The same object feature was associated with reward for blocks of 45-60 trials and then switched to a new rewarded target object feature.

Experimental sessions used the task with two conditions that used either low visual interference and deterministic reward, or high visual interference and probabilistic reward. The three objects either varied in features of one feature dimensions (e.g. different colors), or in features of two dimensions (e.g. different colors and object shapes) constituting low or high visual interference. In the low interference condition, the reward feedback was deterministic (each correct choice resulted in valid feedback), while in the high interference condition the reward feedback was provided with p.085 probability.

From each session we extracted the trials subjects performed during the 20 min. neurochemical sampling event. Analysis calculated the number of completed trials, the total number of reward pulses that the animal received, the performance of the animal (the rate of selecting the highest reward probability object), the rate of reward pulse acquisition (over trials), the average inter-trial interval, reaction times (RTs), and a metric measuring learning speed defined as the first trial in a sequence of 10 trials where performance reaches 70% or higher. Performance of the tasks differed (**Fig. S6A**). In sessions using the task with high interference and probabilistic reward subjects learned the feature-reward rule slower (Wilcoxon’s rank sum, p = 0.0001), showed lower plateau accuracy when learning was completed (Wilcoxon’s rank sum, p = 0.0001), received less total number of rewards, (Wilcoxon’s rank sum, p = 0.0001) and showed slower reaction times on error trials (Wilcoxon’s rank sum, p = 0.0110) (**Fig. S6A**). Experimenters provided less reward pulses for the easier task to keep the total reward similar, which accounts for the lower reward rate in easy task difficulty since the number of trials completed are consistent across both difficulty levels.

#### Measuring heart rate and oxygen saturation levels

While the animals performed the behavioral task and were undergoing neurochemical sampling, we collected heart rate and SpO_2_ data at 0.25 Hz sampling rate using a pulse oximeter (PalmSAT 2500, Nonin Inc, MN) clipped to the ear lobe of the subjects. Overall, we did not observe apparent correlations with the exception of acetyl-cholinergic levels in the striatum showing anti-correlations with both, heart rate (r = -0.475, p = 0.0059) and oxygen levels (r = -0.364, p = 0.0268) (**Fig. S7A**).

### Statistical Analysis

#### Analysis of total concentrations of neurochemicals per area

We analyzed individual concentration values per region, monkey and rest or task state on a logarithmic scale used Kruskal Wallis tests for multiple comparisons with post hoc Wilcoxon’s rank sum for paired test, and FDR corrected per area (**Fig. 2A, 3A, S1**). Across analyses we defined outlier datapoints as those exceeding 2 standard deviations and excluded 7/6/8/4/5 of outlier datapoints in the PFC for the compounds glutamate/GABA/acetylcholine/choline/serotonin and 6/8/5/6/6/6 of outlier datapoints in the striatum for the compounds glutamate/GABA/acetylcholine/choline/dopamine/serotonin.

#### Analysis of the difference between rest and task states

We quantified how neurochemical concentrations changed between rest and task states by collecting non-NaN rest-task pairs (rest-easy or rest-hard) from each experimental session per neurochemical, monkey, area (**Fig. 2B, 3B**). After removing outliers (std (2) <), we converted the raw concentrations into z scores centered around the geometric mean within every session using the following formula:

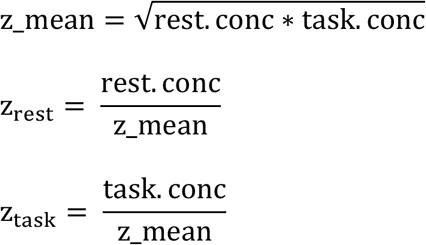

This allowed us to standardize within each session and derive meaningful ratios for both rest and task concentrations. We then tested whether the rest versus task state difference for each neurochemical deviated from zero using Wilcoxon’s sign rank test. To test whether the rest versus task state difference between the neurochemicals differed significantly, we used Kruskal Wallis test. To find which specific neurochemical standard difference differed significantly, we used post hoc Wilcoxon’s rank sum test with FDR correction for multiple comparisons.

#### Decoding cognitive state using mixed effects logistic regression models

We analyzed how combinations of neurochemicals distinguished the cognitive task state from the rest state using mixed logistic regression modeling (**Fig. 2C, 3C, S3**). Neurochemical concentrations (predictors) were organized column-wise with their outcome variables (rest or task) session-wise. To avoid dealing with missing data, we removed sessions with missing values for neurochemicals. After removal, we had 28 sessions, or 56 samples for the PFC, and 23 sessions, or 46 samples in the striatum. Both datasets were balanced (chance = 50%). We then converted raw concentrations into z-scores centered around the geometric mean. Because logistic regression is sensitive to outliers, this standardization allowed us to compress the outliers into ratios and derive meaningful scores during both rest and task. We then found all possible combination of predictors and individual predictors (PFC: 31, Striatum: 63) and ran mixed effects logistic regression with 7-fold cross-validation for every individual and combination of predictors. Sessions were randomly assigned to each fold to reduce variance from small sample size. For our overall model, we used monkey and sessions as random effects, and for individual monkey models, we considered only sessions as our random effect. Laplace approximation was used to derive distribution of random effects to reach an approximate maximum likelihood (Vonesh, 1996). During cross-validation, predictions were made at the population level (marginal predictions) without subject specific random-effects (Colby and Bair, 2013). We used Student’s t test to understand if the accuracies of the individual models were significantly different from chance (50%). Accuracy is the number of correct predictions (true positives + true negatives) divided by the total predictions made. Error estimates represent standard error of the mean across folds.

#### Analysis of correlations among neurochemical levels and performance/autonomic metrics

To understand how neurochemicals co-vary during rest or task state, we sorted rest concentrations and task concentrations separately into 3 matrices: one for each monkey and a pooled matrix. Due to the non-parametric distribution of the neurochemicals, we used the robust Spearman’s partial rank correlation test. We performed Spearman’s partial rank correlation test in a similar manner to control for the linear effects of other neurochemicals on the pair (**Fig. 2D,E, 3D,E**). We applied Benjamini-Hochberg false discovery rate correction to these p-values with an alpha of 0.05.

## Acknowledgments

This work was supported by the National Institute of Mental Health R01MH129641 (TW). The funders had no role in study design, data collection and analysis, the decision to publish, or the preparation of this manuscript.

## Author contributions

Conceptualization: SAH, AN, JP, TW

Methodology: SS, SAH, AN, SL, KSR, JP, TW

Investigation: SAH, AN, SL, KSR

Visualization: SS, SL, TW

Supervision: JP, TW

Writing—original draft: SS, TW

Writing—review & editing: SS, SAH, AN, SL, JP, TW

## Competing interests

All other authors declare they have no competing interests.

## Data and materials availability

Data and code will be available at the time of publication at an open-access repository.

## Notes

### Competing Interest Statement

The authors have declared no competing interest.

