## Supplementary Material for "Cognitive engagement induces area-specific fingerprints of dopamine, acetylcholine, serotonin, glutamate and GABA in prefrontal cortex and striatum"

**This PDF file includes:**

- Supplementary Text
- Figures. S1 to S8
- Tables S1 to S2
- Supplementary References

**Supplementary Text**

**Chemicals, reagents and materials.** The LC-MS-grade solvents methanol (MeOH), acetonitrile (ACN), isopropanol (IPA) and water, as well as the acetylcholinesterase inhibitor phenylmethylsulfonyl fluoride (PMSF) were purchased from Fisher Scientifc. Formic acid, polyacrylonitrile (PAN), N,N-dimethylformamide (DMF) and the standards of neurotransmitters: γ-aminobutyric acid (GABA), glutamic acid (Glu), acetylcholine (ACh), histamine (Hist), serotonin (5-HT), dopamine (DA) and choline (Cho) as well as their deuterated analogues were purchased from Millipore Sigma (Oakville, ON, Canada). Epinephrine (Epi), norepinephrine (NE) and their deuterated analogues were obtained from Cerilliant Corporation (Round Rock, TX, USA). Choline-D9, was purchased from Cambridge Isotope Laboratories (Tewksbury, MA, USA). The reagents used for synthesis of hydrophilic-lipophilic balance polymer particles functionalized with strong cation exchange groups, as well as compounds for preparation of PBS were purchased from Millipore-Sigma. The stainless steel wire (stainless steel grade AISI 304, 150 µm diameter) used for manufacturing of SPME probes was purchased from Unimed S.A. (Lausanne, Switzerland). The stainless steel tubing used as guiding cannulas (270 µm O.D.; 200±5 µm I.D.) was obtained from Vita Needle (Needham, MA, USA).

**Quantitation of neuromodulators.** Individual stock standard solutions of all targeted neuromodulators were prepared in methanol or water with 0.1% formic acid at a concentration of 1 mg/mL and stored at −80 °C for maximum of one month. In order to calculate the amounts of neuromodulators extracted by each probe, calibrator standards prepared in the same desorption solvent as real samples were analyzed in the same batch. This instrumental calibration curve was prepared in the range of 0.1-200 ng/mL by a serial dilution of the stock standard mixture of all compounds at 1µg/mL. The IS concentration was kept constant at 20 ng/mL in all calibrators, identically as in the real samples. The amounts extracted were calculated based on linear regression equation obtained from the analytical signal (the ratio of chromatographic peak areas of analytes and their corresponding IS) plotted against the concentration.

In order to calculate the concentrations of neuromodulators sampled from brain tissue, matrix-matched external calibration approach was used. The surrogate matrix consisted of 2% agar gel mixed with brain homogenate in the ratio 1:1 (*v/w)*. The homogenized brain tissue was earlier incubated with 1 mM PMSF for 1 h at 37°C to prevent enzymatic digestion of acetylcholine in the calibrator samples. Due to several target compounds being present in brain homogenate at high concentrations (e.g. for glutamate and choline the "blank" brain homogenate matrix doesn't exist), their quantitation was based on signals of their deuterated isotopologues. The calibrator samples were prepared in the surrogate matrix with concentrations of neuromodulators ranging from 5 to 3000 µg/mL for the isotopically labelled compounds or from 10 to 2000 ng/mL for the remaining compounds. The extractions were carried out with SPME probes manufactured and pre-treated identically to the probes used for *in vivo* sampling and using the same 20 min extraction time and desorption conditions as for the real samples. The amounts of neuromodulators extracted from the calibrator samples were determined in the same way as described above and plotted against concentrations of calibrators. The resulting weighted linear regression equations were applied to the amounts of neuromodulators extracted from the *in vivo* samples, yielding values of concentrations of the compounds of interest in brain.

Quantitation of neurochemicals derivatized with benzoyl chloride was conducted in the same way as described above, by aliquoting 20 uL of the calibration curve and in vivo extracts, evaporating to dryness at 30°C, adding 10µL of 50 mM sodium tetraborate buffer (pH 9.2) and subsequent addition of 10 µL of benzoyl chloride solution (2% in acetonitrile, *v/v*).

The limits of detection (LOD) were estimated as the levels corresponding to the signal to noise ratio of 3 and were calculated based on the signal of blank calibrator sample (considered as the noise).

**Post-desorption derivatization with benzoyl chloride**. One of the challenges in quantitative analysis of low-concentration neurotransmitters via LC-MS is the small size of the target molecules, placing them in the most highly populated region of the MS spectrum, giving rise to interferences that result in increased noise and decreased signal-to-noise (S/N) ratios (Renaud et al., 2017). The post-desorption derivatization with benzoyl chloride was pursued to decrease the MS background and interferences by increasing the target analytes' hydrophobicity and m/z values which in turn reflects the increase in the S/N ratio of the analytes. Significant improvements in LOQ concentrations were achieved in this way for serotonin (from 50 ng/g to 10 ng/g after derivatization) and GABA (from 5000 ng/g to 125 ng/g after derivatization). Even though acetylcholine does not undergo benzoyl chloride derivatization due to its quaternary ammonium group, LOQ for this analyte has also been significantly improved (from 50 ng/g to 10 ng/g) likely as it benefits from different LC-MS conditions employed for the derivatized analytes.

**Properties of SPME extraction that make it a versatile sampling procedure.** The use of SPME to sample neurochemicals in brain tissue in vivo has been validated in prior in-vitro and in-vivo studies (Hassani et al., 2019; Lendor et al., 2019a; Lendor et al., 2019b; Bogusiewicz et al., 2021; Hassani et al., 2023). To explicitly describe the qualities of SPME and how they relate to existing approaches we compare the SPME approach with microdialysis (MD) and cyclic voltammetry in **Table S2**. SPME partly complements or outperforms these alternative techniques along various domains, with several practical considerations of the SPME approach to in-vivo brain sampling as outlined below:

1) ***Spatial Resolution.*** In principle, the spatial resolution of the SPME measurements is capable of reflecting laminar concentration gradients across areas as small as several tens of micrometers. With a probe placement traversing compartments of differing concentrations, these gradients can be imprinted on the SPME device, preserving this spatial distribution information until analysis. However, due to the challenges related to instrumental sensitivity for neurotransmitter measurements related to their size and hydrophilicity, as outlined above, in our protocol we employed a 3mm length of coating the probe with a 50 µm of extracting phase thickness and optimized placing the probe coating in the grey matter using electrophysiological recordings of neural activity as guide (*see* Methods). The absolute spatial resolution of SPME is difficult to estimate as SPME operates through diffusion and dynamic equilibration similar to MD. Moreover, due to the MD and SPME probe size ranges, the sampled compartment includes predominantly the extracellular space, with some contributions from the vascular system and neural cells ruptured upon probe insertion  As a consequence, the volume of the sampled compartment will depend on sampling time, diffusion coefficient of the target analytes, as well as the rate of diffusion within the tissue (Sykova, 2004). The versatility of the SPME approach allows for customization of the coated probe length, allowing it to be adjusted to the size of the sampled brain area.
2) ***Temporal Resolution.*** The amount of analyte extracted as a function of extraction time (the extraction time profile) reflects the kinetics of SPME sampling and it can aid empirical selection of extraction time for a particular type of experiment. While both pre-equilibrium and equilibrium extraction yield quantitative results, one of the challenges of in-vivo brain analysis is the dynamic character of the release and uptake of each of the neurochemicals. The specificity of theses dynamics effectively results in multiple equilibration profiles throughout the sampling event (Zhang et al., 2010; Lendor et al., 2019b), because the amount of an analyte extracted via SPME is proportional to its average concentration in the target brain area over the duration of the measurement (Poole et al., 2016). Employing short extraction times enhances the temporal resolution of the sampling, however in practicality, the dynamic neurochemical events such as neurotransmitter release, degradation, and reuptake occur on a sub-second timescale, well below the typical SPME or MD sampling times of several or several tens of minutes. Techniques such as SPME or MD are however suitable for studies with behavioral components as shown by this study.
3) ***Utility and Ease of Use***. SPME devices are highly amenable to modifications according to the scope and requirements of the study. The extracting phase thickness and length are particularly

important tunable features, as in order to represent the brain's undisturbed chemistry at the time of sampling, non-depletive extraction conditions are necessary. SPME fulfills these requirements with the addition of biocompatibility (Kasperkiewicz et al., 2023). SPME lacks MD's capabilities of continuous sampling and on-line coupling to detection devices, however this presents the upside of eliminating the risks of adverse biofouling effects common to biomedical devices (Harding and Reynolds, 2014), as the transient character of SPME sampling and probe insertion does not require long-term implantation. Similarities in size and format between SPME and recording microelectrodes used for electrophysiology reduce the adjustment requirements and allow for adapting SPME measurements to existing microelectrode positioning systems. Additionally, SPME can take advantage and build upon the prevalence of MD in neuroscience, as the probe positioning interface can be built by modifying a MD setup. This also applies to post-processing and detection methods, as instrumental analysis of samples collected via SPME and MD are treated similarly, with the main difference being the calibration strategies to achieve quantitative results (Cudjoe et al., 2012).

4) ***Extension of SPME measurements beyond classical neuromodulators.*** Operating as a *chemical biopsy* device with no tissue collection or removal, SPME offers diverse fit-for-purpose modes for *in vivo* analysis. Full customization of extracting phase chemistries and sizes contributed to recent expansion of monitoring and quantitation of various chemicals in brain in vivo. In recent years, in vivo SPME brain sampling has been applied to neurologically relevant targets including endocannabinoids (Oliveira et al., 2026), oxylipins (Napylov et al., 2020), nicotine metabolites (Zhang et al., 2025) or xenobiotics donepezil (Hassani et al., 2023) and ketamine (Lendor, 2020), as well as non-targeted studies investigating metabolomic and lipidomic changes in the brain after various interventions (Boyaci et al., 2020; Lendor et al., 2020).

**Relationship of acetylcholine and choline levels in striatal and cortical circuits.** We found that acetylcholine (ACh) and choline (Cho) showed no apparent correlation in the PFC at rest and a moderate and statistically non-significant correlation during task engagement ($r=0.22$, **Fig. 2D-F**), while striatal ACh-Cho correlations similarly were low (rest: $r=0.2$; task: $r=0.07$). Similarly, ACh and Cho were not apparently differently modulated by the more difficult than the easier task condition in the PFC (**Fig. S5**) or the striatum (**Fig. S8**). Furthermore, the decoding model showed ACh and Cho modulated inversely with task engagement by contributing to the prediction of task state with moderate increases of ACh and moderate decreases of Cho levels in the PFC (**Fig. 2C**) and in the striatum (**Fig. 3C**).

To understand this relationship further, we compared the ratio of ACh/Cho in the rest and task states and found that the ratio increased in the PFC (Wilcoxon's test, $p = 0.003$) and at a statistical trend also in the striatum (Wilcoxon's test, n.s. $p = 0.0885$) (**Fig. S8**). While extracellular ACh and Cho don't covary in either state, the balance between the two changes with cognitive engagement. As our decoding results show (**Fig. 2C & 3C**), this change in balance is due to increased extracellular acetylcholine during the task state. Interestingly, in the PFC, where the ACh/Cho relationship distinguishes rest and task, ACh is mainly covarying with glutamate across both states. In the striatum, where the distinction between the ratio is weaker between rest and task, Cho mainly covaries with glutamate. This may reflect a region-specific difference in ACh/Cho balance maintenance.

ACh and Cho share a unique relationship as Cho is both the precursor and metabolite of ACh. This makes it difficult to interpret cholinergic transmission using Cho levels. Some studies suggest a reciprocal relationship between the two, such as when extracellular ACh increases, Cho decreases, so Cho may be informative of cholinergic transmission (Koshimura et al., 1990; Ikarashi et al., 1997). Other studies suggest Ach-derived Cho is rapidly cleared, therefore increases in extracellular Cho levels may not be informative of cholinergic transmission (Parikh et al., 2004; Sarter and Parikh, 2005). In our study, we find in the PFC and moderately in striatum that ACh and Cho levels – measured in a 20 min sampling event - don't covary within a state due to a balance likely maintained by factors such as density of low vs high affinity transporters, CHT regulation in the region, or diffusion of plasma Cho (Atweh et al., 1975; Simon et al., 1976; Sarter and Parikh, 2005). With task engagement, extracellular ACh transiently increases, shifting the balance between ACh/Cho (Dalley et al., 2001; Arnold et al., 2002). Together, these considerations suggest that ACh/Cho balance, rather than acetylcholine alone, serves as a region-specific indicator of cholinergic tone, and adapts dynamically based on cognitive demand.

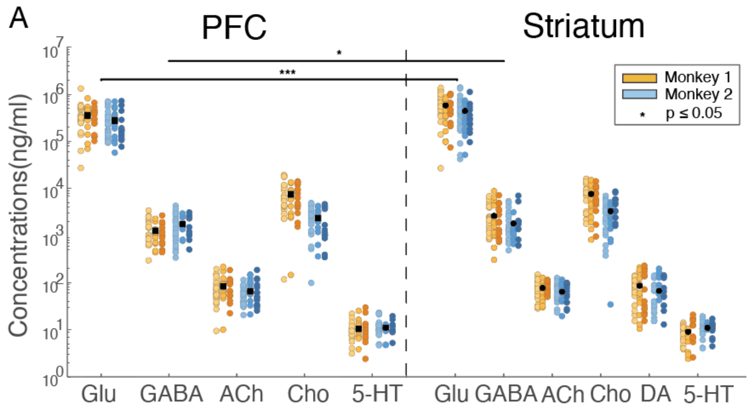


B PFC

| | GABA | ACh | Cho | 5-HT |
|---|---|---|---|---|
| Glu | 1.41e-38 | 4.52e-39 | 6.59e-39 | 1.04e-35 |
| GABA | | 9.75e-39 | 2.36e-13 | 1.97e-35 |
| ACh | | | 2.74e-38 | 1.64e-33 |
| Cho | | | | 1.04e-35 |

C Striatum

| | GABA | ACh | Cho | DA | 5-HT |
|---|---|---|---|---|---|
| Glu | 4.53e-38 | 2.07e-38 | 9.64e-39 | 6.7e-36 | 2.56e-35 |
| GABA | | 1.35e-37 | 9.72e-11 | 3.54e-35 | 1.24e-34 |
| ACh | | | 4.18e-37 | 0.8715 | 8.28e-35 |
| Cho | | | | 6.76e-35 | 3.49e-35 |
| DA | | | | | 1.56e-30 |

**Fig. S1. Statistical comparison of neurochemical concentration level sin POFC and striatum.** (**A**) Neurochemical concentrations measured in the PFC (*left*) and striatum (*right*) across all sessions of subject 1 (*yellow*) and 2 (*blue*). Concentrations in the PFC and striatum differed for Glutamate (Wilcoxon's test, FDR corrected, p=3.48e-05), and GABA (Wilcoxon's test, FDR corrected, p=1.97e-02). (**B**,**C**) Raw concentration values differed for all pairs of neurochemicals: p-values from post-hoc Wilcoxon's tests comparing total neurochemical concentrations in PFC (*B*) and striatum (*C*).

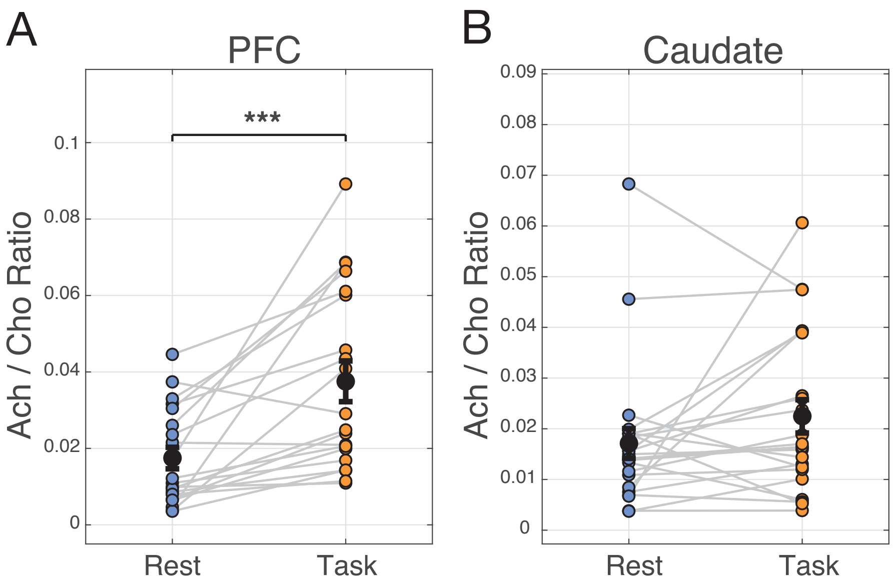


**Fig. S2. Ach-Cho balance changes with cognitive engagement.** (**A**) Ratio of Ach/Cho in the PFC in Rest versus Task state (Wilcoxon's test, p = 0.0003), and (**B**) ratio of Ach/ Cho in the caudate in Rest versus Task state (n.s. p = 0.0885. Black dots are means with SE error bars. *** stars above horizontal bars denote p<0.001

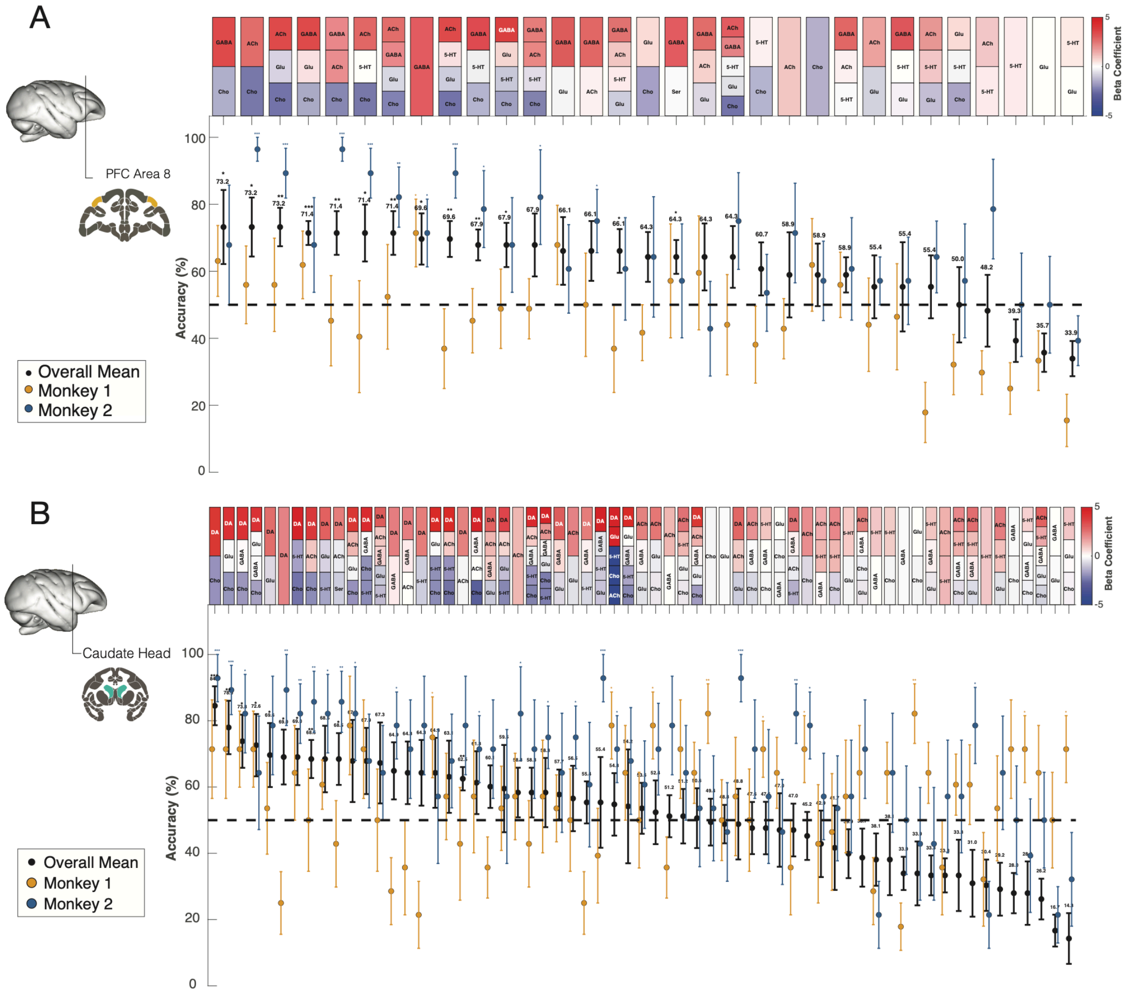


**Fig. S3. Predicting the Cognitive vs Rest state with combinations of neurochemical concentrations using mixed logistic regression models** (**A**) Accuracy of classifying Task vs Rest state (best ranked on the left) from of all combinations of neurochemicals sample in the PFC using a generalized non-liner mixed effects model with 7-fold cross-validation. The top row shows the color-coded beta-coefficients of the neurochemicals underlying the accuracy shown in the bottom panel. Red colors denote neurochemicals predicting the task state with increased concentrations relative to the rest state; blue denotes neurochemicals with levels reduced in the task state. (**B**) Same as A for the neurochemicals sampled in the striatum. For *A* and *B* the first ten best ranked neurochemical combinations are also shown in *Fig. 2C* and *Fig. 3C*, respectively, and shown here to facilitate comparison across all neurochemicals.

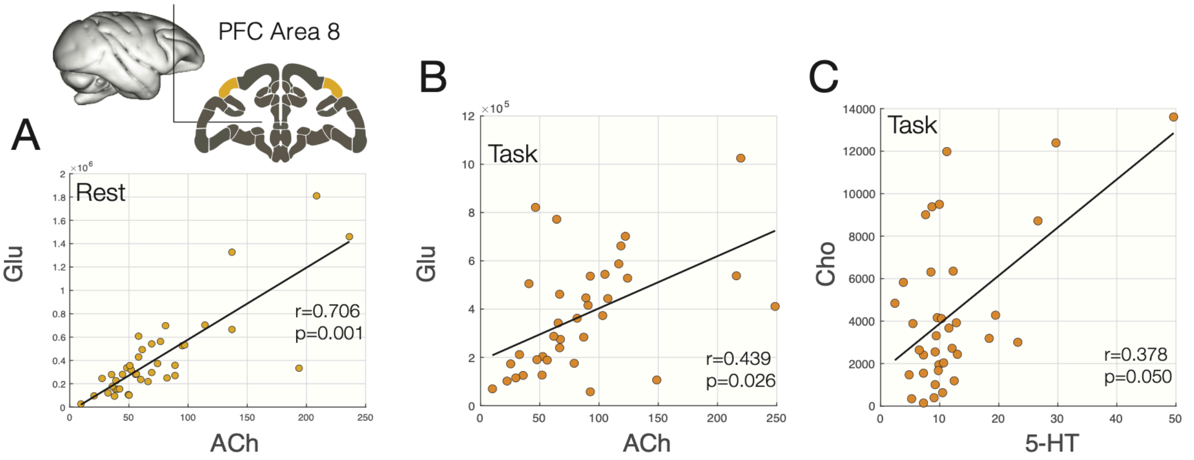


**Fig. S4. Correlations of neurochemical concentration of glutamate, acetylcholine and serotonin in the prefrontal cortex.** (**A**) Spearman's partial rank correlation (r) of Glu and ACh when subjects were at rest (r = 0.706, p = 0.001), and (**B**) when subjects engaged in the task (r = 0.439, p = 0.026). (**C**) Spearman's partial rank correlation between Cho and Ser during cognitive task engagement (Spearman's partial rank correlation: r=0.378, p=0.05; Monkey 1: r=0.63, p<0.05; Monkey 2; r=0.62, p<0.05).

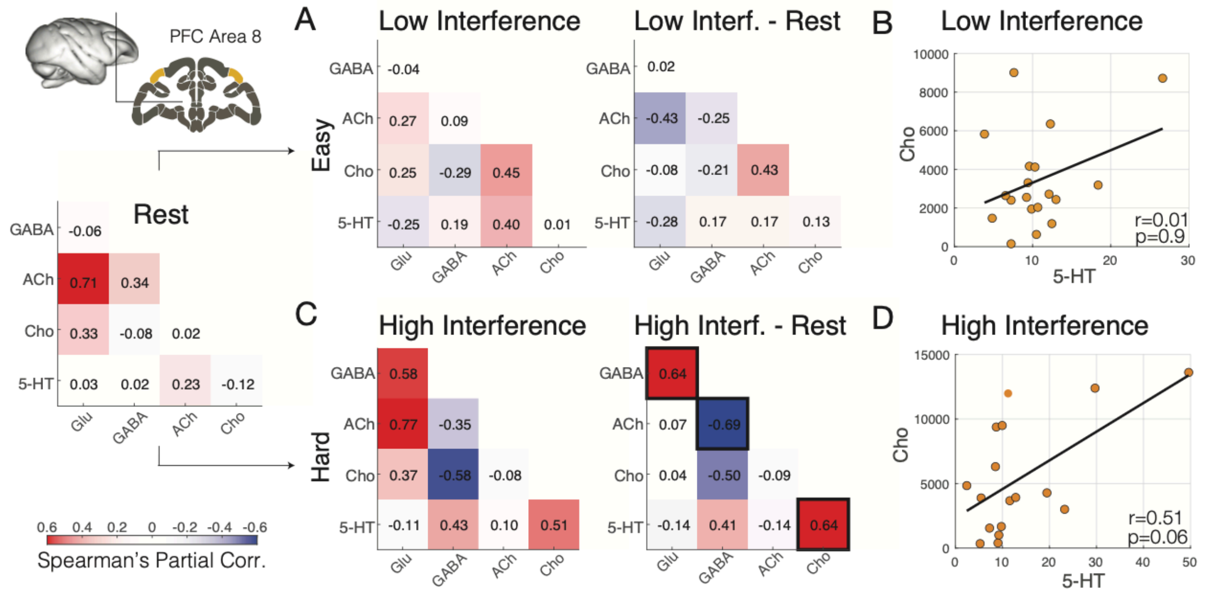


**Fig. S5. Neurochemical co-modulation in the PFC at rest and during cognitive engagement in task conditions with low and high interference** (**A**) Spearman's partial rank correlations of neurochemicals during rest (*leftmost*), and during the low-interference task condition (*left*), and the correlation difference [Low interf. task minus Rest] (*right*). (**B**) Correlation of choline (*y-axis*) and serotonin (*x-axis*) during performance of the task with low interference. (**C,D**) Same as A,B, for neurochemical levels during cognitive engagement in the task condition with high interference. Cells with thick outline denote differences at $p<0.05$.

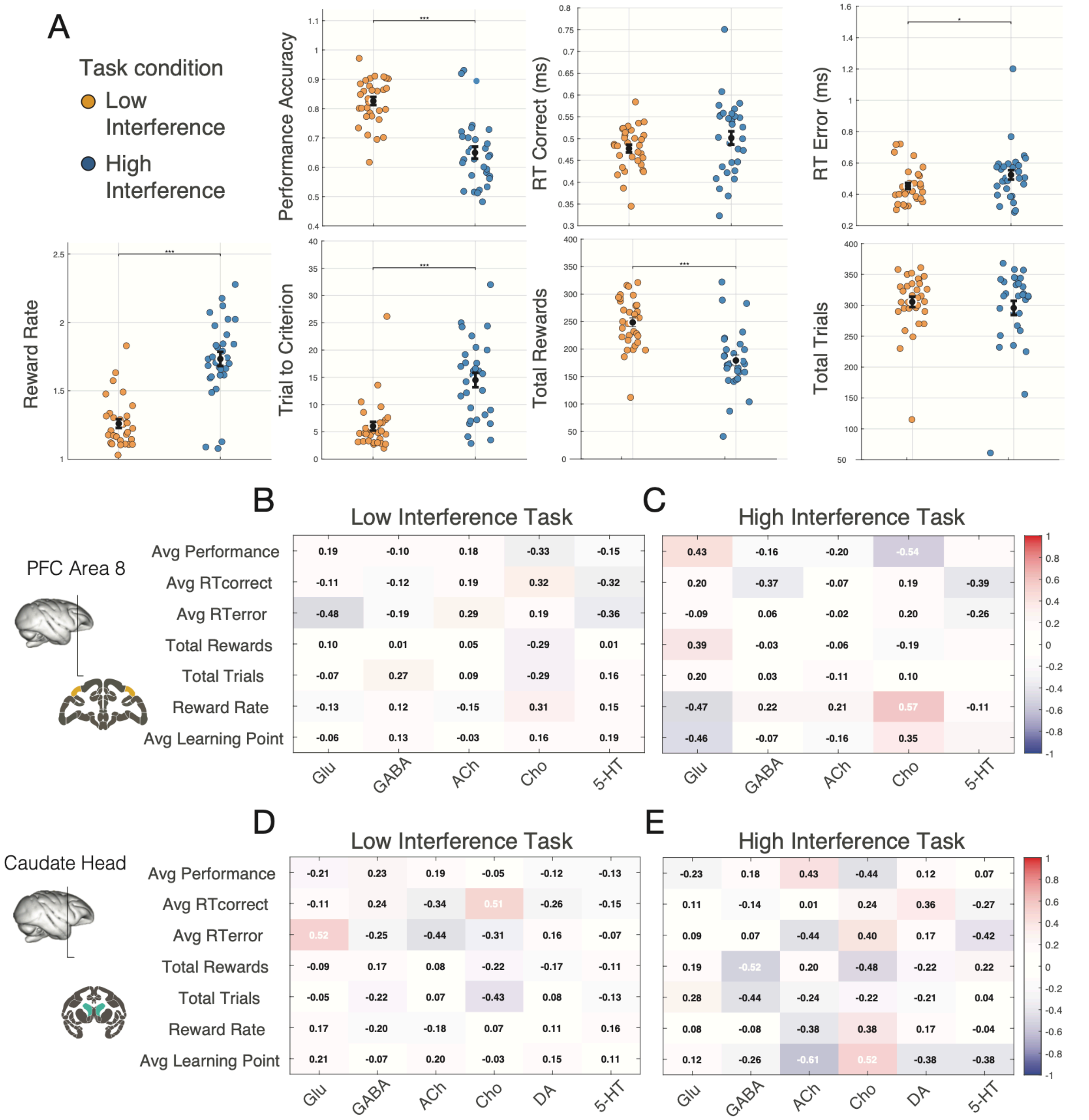


**Fig. S6 Relationship of neurochemical level and the cognitive demands on interference control.** (**A**) Seven performance metrics calculated for all trials obtained during the 20 min. neurochemical sampling periods shown across sessions for the low (*yellow*) and high (*blue*) interference task conditions. Black dots are means with SE error bars. * and *** stars above horizontal bars denote $p<0.05$ and $p<0.001$. While the total number of trials performed during the 20 min SPME sampling was similar between task conditions, all major performance measures differed between task conditions. (**B**,**C**) For the PFC, correlations of the neurochemical levels (*x-axis*) and the performance metrics (*y-axis*) in the task condition with low interference (*B*) and high interference (*C*). (**D**,**E**) Same as *B*,*C* for the anterior striatum. There was no correlation in *B-E* passing the $p < 0.05$ criterion.

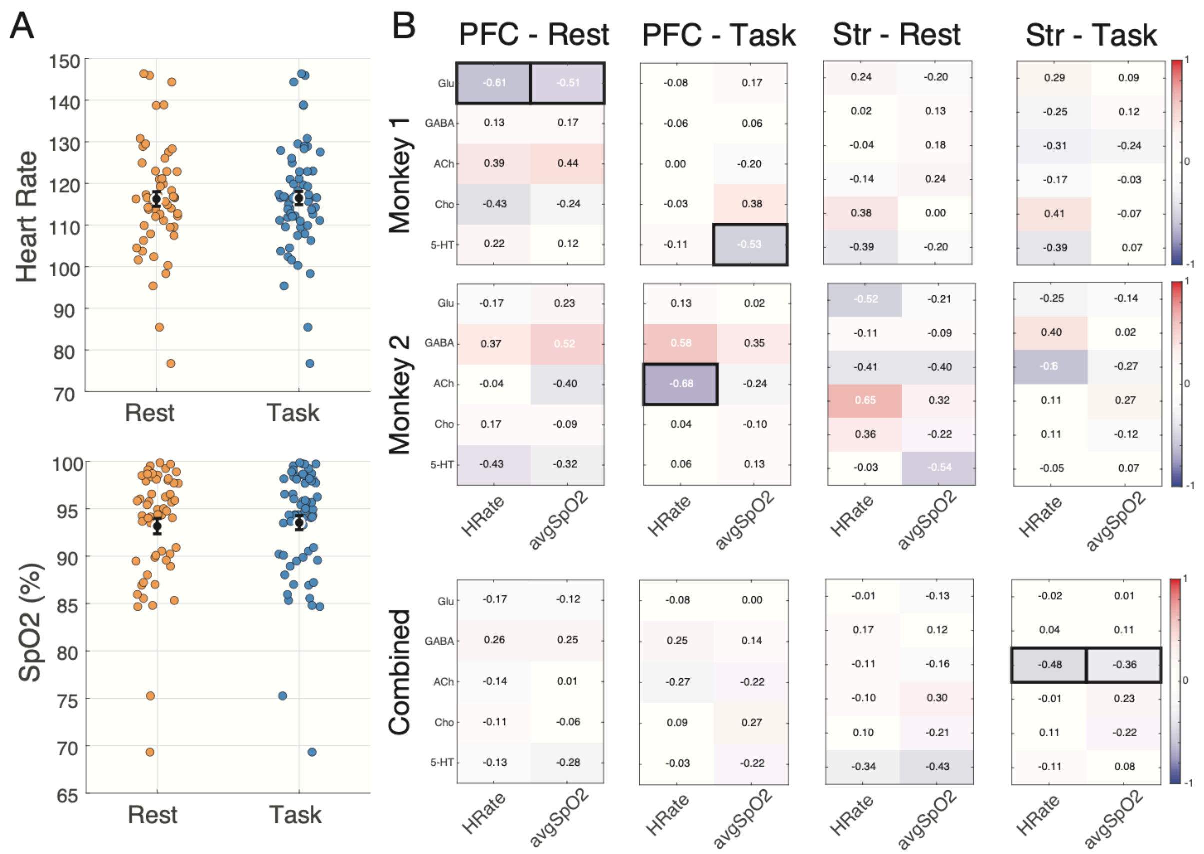


**Fig. S7. Correlation of neurochemical concentrations, heart rate and $SPO_2$ levels.** (**A**) Average heart rate (*top panel*) and $SPO_2$ levels (*bottom panel*) measured during the 20 min sampling at rest (*yellow*) and during task performance (*blue*) across sessions. (**B**) Correlations of heart rate and $SPO_2$ levels (*x-axis*) with of neurochemical levels (*y-axis*) in the PFC at rest, during the task (*two leftmost columns*) and in the striatum at rest, during the task (*two rightmost columns*). Top two rows are from sessions of subjects 1 and 2, bottom row shows correlations combined across sessions from both subjects. Thick outlined cells denote correlations at $p<0.05$. Across both monkeys combined, striatal acetylcholine correlated with heart rate ($r = -0.475$, $p = 0.0059$) and $SPO_2$ levels ($r = -0.364$, $p = 0.0268$).

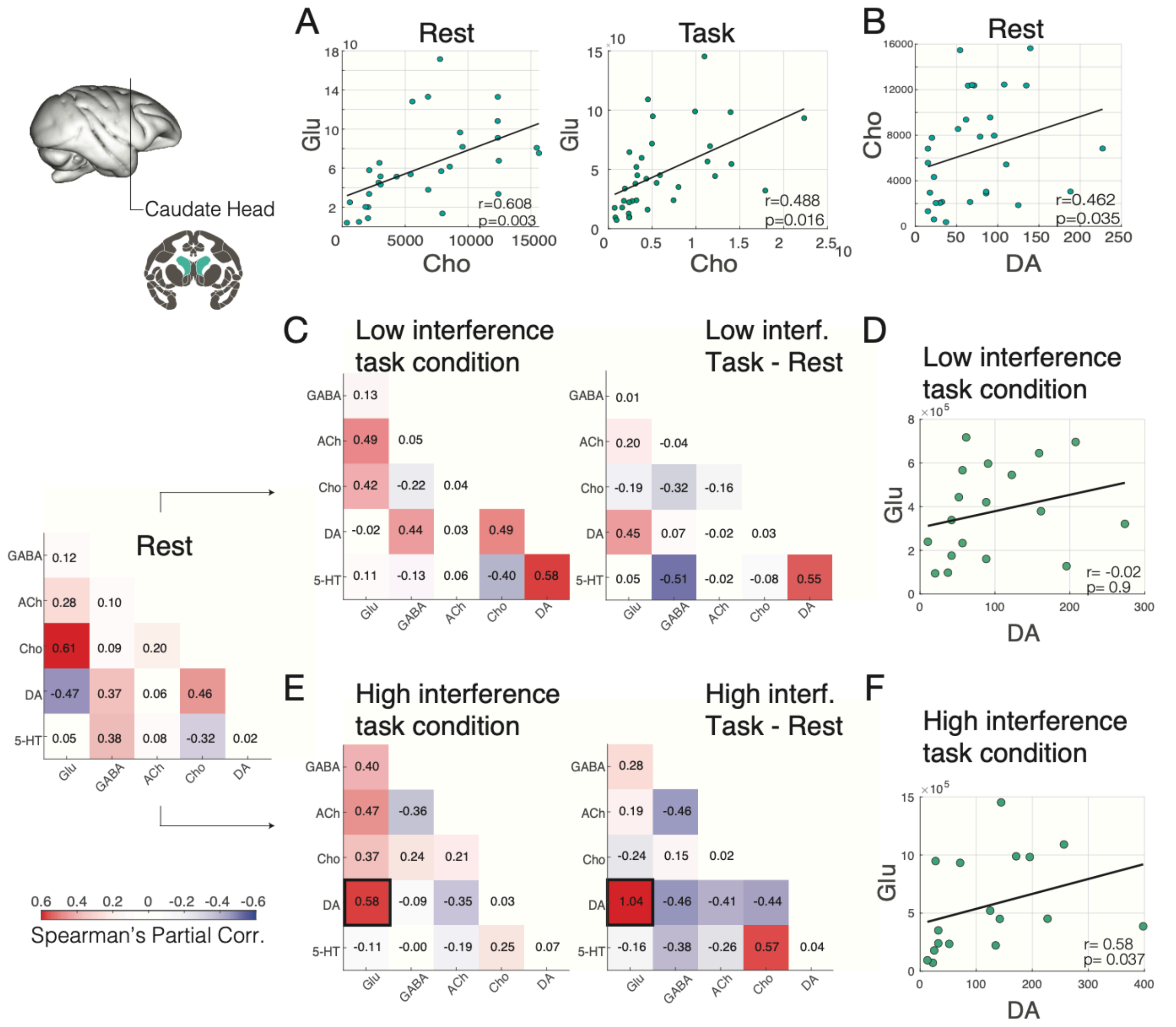


**Fig. S8. Neurochemical co-modulation in the striatum at rest and during cognitive engagement in task conditions with low and high interference.** (**A**) Spearman's partial rank correlation between Glu and Cho during the Rest state (*left:* r = 0.608, p = 0.003) and Task state (*right*: r = 0.488, p = 0.016) and (**B**) between Cho and Dop are at rest (r = 0.462, p = 0.035). (**C**) Spearman's partial rank correlation of neurochemicals during the low-interference task condition (*left*), and the correlation difference (*right*) of the [Low interf. task condition minus Rest (*see leftmost panel*)]. (**D**) Spearman's partial rank correlation of Glu (*y-axis*) and Dop (*x-axis*) during performance of the task with low interference. (**E**,**F**) Same as *C*,*D* for neurochemical levels during cognitive engagement in the task condition with high interference. Cells with thick outlines denote differences at p<0.05.

### Supplementary Tables

**Table S1**. Details of the LC-MS/MS methods employed for quantitation of targeted neurochemicals

| | | No derivatization | Benzoyl Chloride Derivatization |
|---|---|---|---|
| Tandem Mass Spectrometry | | **Thermo Quantiva Triple Stage Quadrupole** (heated electrospray ionization source; **positive ion mode**) | |
| | Analytes (IS) | Dopamine<br>Choline<br>Glutamate | Serotonin<br>Acetylcholine<br>GABA |
| | Spray voltage [kV] | 3.5 | |
| | Sheath gas [Arb] | 45 | |
| | Auxiliary gas [Arb] | 13 | |
| | Sweep gas [Arb] | 1 | |
| | Ion transfer tube temperature [°C] | 342 | |
| | Vaporizer temperature [°C] | 358 | |
| | Acquisition mode | SRM, dwell time 50 ms | |
| | Data acquisition and processing software | Xcalibur 4.0, Trace Finder 4.1 | |
| | Monitored transitions | Dopamine 154 → 91<br>Dopamin-D4 158 → 141<br>Choline 104 → 60<br>Choline-D9 113 → 69<br>Glutamate 148 → 84<br>Glutamate-D5 153 → 88 | BzCl-Serotonin 385 → 105<br>BzCl-Serotonin-D4 389 → 105<br>Acetylcholine 146 → 87<br>Acetylcholine-D9 155 → 87<br>BzCl-GABA 208 → 105<br>BzCl-GABA-D6 214 → 105 |
| Ultra High Pressure Liquid Chromatography | | **Dionex UltiMate 3000 UHPLC** | |
| | Column | Phenomenex Kinetex PFP, 1.7 µm, 2.1 x 100 mm | Thermo Scientific Hypersil GOLD C18, 1.9 µm, 2.1 x 50 mm |
| | Mobile phase A | $H_2O$/MeOH/ACN 90:5:5 + 0.1% FA | $H_2O$ + 0.1% FA + 3.5 mM ammonium formate |
| | Mobile phase B | ACN/ $H_2O$ 90/10 + 0.1% FA | MeOH + 0.1% FA + 3.5 mM ammonium formate |
| | Flow rate [µL/min] | 400 | 400 |
| | Column temperature [°C] | 35 | 30 |
| | Samples temperature [°C] | 5 | 5 |
| | Injection volume [µL] | 10 | 10 |
| | Gradient [%B] | 0 min-100%; 1 min-100%; 4 min-0%; 5.5 min-0%; 5.6 min-100%; 6.5min-100% | 0 min-0%; 1 min-0%; 4min-100%; 5 min-100%; 5.1 min-0%; 6min-0% |

**Table S2.** Comparison of methods capable of measuring single or multiple neurochemicals in vivo. Temporal resolution, spatial resolution, sensitivity, capability to measure neuro-active and non-neuro-active compounds, in vivo feasibility and cost. PET: positron emission tomography; NHP: non-human primate; MS: mass spectrometer.

| | PET imaging | Electro-chemistry | Fluorescent Biosensors | Micro-dialysis | Solid phase micro-extraction |
|---|---|---|---|---|---|
| *Temporal Resolution* | Minutes | Highest (millisecond range) | Highest (millisecond range), but risk photobleaching at long durations | 1-30 minutes; dependent on MS sensitivity, target etc. | <5-30 minutes; dependent on MS sensitivity, coating thickness, target etc. |
| *Spatial Resolution* | Voxel | High; surface area may vary (relevant for enzyme-based methods) | High | Diffusion based; surface area may vary | Diffusion based; surface area may vary |
| *Sensitivity* | Indirect measurement via competitive radiolabeled species | High | High, but concentration values are relative per individual | Depending on post-hoc methods (i.e. MS) | Depending on post-hoc methods (i.e. MS) |
| *Neuro-active targets* | A few at most | A few at most | Depends on biosensor availability for the organism | Many | Many |
| *Non-neuro-active targets* | No | No | No | Yes, greater efficacy for hydrophilic compounds | Yes, greater potential efficacy for hydrophobic compounds |
| *In vivo feasibility* | Difficult in awake, behaving animal models; movement highly restricted | Good (low reliability in NHPs) | Good, often requires chronic implant of fiber | Good, often requires chronic implant of cannula for repeated measurements | Very good; robust placement of multiple simultaneous probes and repeatable acute measurements |
| *Cost* | High | Requires special equipment | Requires special equipment | Requires special equipment | Easy to port to an acute micro-electrode setup; requires a chemistry core |
